# PathRacer: racing profile HMM paths on assembly graph

**DOI:** 10.1101/562579

**Authors:** Alexander Shlemov, Anton Korobeynikov

## Abstract

Recently large databases containing profile Hidden Markov Models (pHMMs) emerged. These pHMMs may represent the sequences of antibiotic resistance genes, or allelic variations amongst highly conserved housekeeping genes used for strain typing, etc. The typical application of such a database includes the alignment of contigs to pHMM hoping that the sequence of gene of interest is located within the single contig. Such a condition is often violated for metagenomes preventing the effective use of such databases.

We present PathRacer — a novel standalone tool that aligns profile HMM directly to the assembly graph (performing the codon translation on fly for amino acid pHMMs). The tool provides the set of most probable paths traversed by a HMM through the whole assembly graph, regardless whether the sequence of interested is encoded on the single contig or scattered across the set of edges, therefore significantly improving the recovery of sequences of interest even from fragmented metagenome assemblies.

**Availability:** http://cab.spbu.ru/software/pathracer/

## 1 Introduction

The recent advances of metagenomics quickly revealed scalability issues arising from the amount of raw sequencing data necessary to describe complex microbial communities. Even more, due to many inherit problems and challenges the assembly of metagenomic data remains a non-trivial task, thus slowing down biological discoveries [13]. Still, in the majority of cases the interest of researchers lies around the recovery of certain important genes this way the use of additional information about these genes might be crucial for the full gene sequence recovery even from very fragmented assemblies.

The typical way to represent the sequence of the particular gene family is via so-called profile Hidden Markov Models [5]. Recently large databases such as Pfam [8] or NCBIfam-AMR [1] containing thousands of pHMMs emerged. These pHMMs may represent the sequences of antibiotic resistance genes, or allelic variations amongst highly conserved housekeeping genes.

The recent releases of Xander [19] and MegaGTA [11] tools opened the possibility of the gene-targeted metagenomics assemblies, where the trained Hidden Markov Model is used to guide the traversal of de Bruijn graph. Such an approach gives obvious advantage over other assembly methods. Still, these tools might be somehow non-trivial to use, for example, Xander forces a user to build both forward and reverse HMMs and MegaGTA requires original sequences the pHMM was built from. All these requirements makes the usage of these tools non-straightforward as one cannot simply download the HMM from the database and use it straight away. Even more, both tools includes their own genome assembler engines essentially ignoring the recent progress made in the field of genome and metagenome assemblies.

PathRacer is a novel standalone tool that aligns profile HMM directly to the assembly graph produced by modern assemblers in standard GFA format^1^. This way PathRacer could utilize all the features and improvements (e.g. hybrid or mate-pair assemblies) provided by state-of-the-art assemblers. Below we describe algorithmic approaches used in PathRacer and showcase them over several datasets.

## 2 Methods

### 2.1 General Definitions

Let *Sequence graph G* be a directed graph where each vertex (called *position*) is labelled by a letter from the given alphabet *Σ*. Thus, each path in *G* corresponds to some string in *Σ^∗^*. Denote by *V* (*G*) and *E*(*G*) the sets of vertices and edges of *G* correspondingly.

*pHMM graph* is a profile HMM in HMMER 3 *Plan 7* format [6 5] on *Σ*. Profile HMM can be viewed as weighted directed graph with vertices called *states* and edges called *transitions*. A particular state could be of one of the following types: I *(insertion)*, M *(match or mismatch)*, D *(deletion)*; in addition to this there are also two special states denoted by Start and End (see Figure 1). I and M states together are called *emissive states* or *emitters*. Each emissive state *E* contains its own emission probabilities 𝓔 = (*E*,⋅) that define a probability distributions on *Σ*. Edges weights denoted as 𝓣(⋅, ⋅) are transition probabilities; in each vertex sum of its outgoing transitions equals to 1. Note that pHMM graph is *almost* acyclic: the only allowed cycles are simple loops from each I state to itself *(I-loops)*. Therefore, pHMM states could be topologically sorted.

**Fig. 1.**
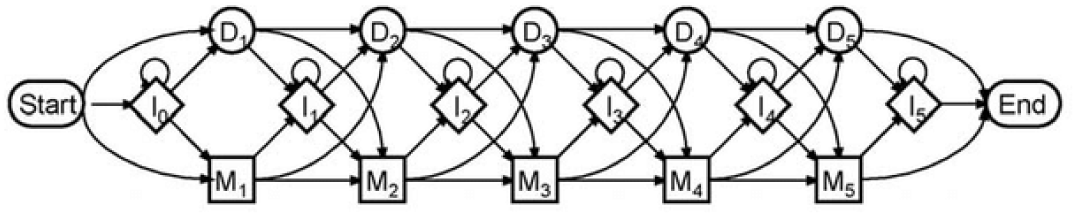
Plan 7 profile hidden Markov model (pHMM) scheme

*Background distribution 𝓑* is a discrete probability distribution on *Σ*. This distribution is an *a priori* distribution of letters in the model. See [6] for more information about background distribution and its role.

For string (*L*_1_*, …, L_n_*) ∈ *Σ^∗^ HMM alignment* is a path in the pHMM graph that contains exactly *n* emitters. If the path begins from Start and ends with End the alignment is called *global*, otherwise the alignment is called *local*. For the sake of simplicity we will consider only the problem of global alignment search. Note that the local alignments could be turned into the global ones by a simple transformation of pHMM graph [6]: insert additional transitions from Start to all emitters and from all emitters to End.

Furthermore, the global alignment is fully defined by the series of its emitters (*E*_1_*, …, E_n_*). Indeed, the “missed” intermediate D states in the alignment path could be unambiguously reconstructed due to a particular structure of pHMM.

For the given pHMM graph, background distribution 𝓑, string 𝓛 = (*L*_1_*, … L_n_*), and its global alignment 𝓐 = (*E*_1_*, …, E_n_*) alignment score *Score*(*𝓛, 𝓐*) is defined as:

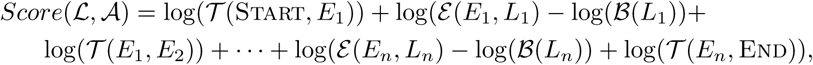

where 𝓣(*E_i_, E_i_*_+1_) may include intermediate D states 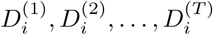:

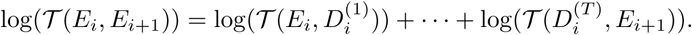

This score is essentially

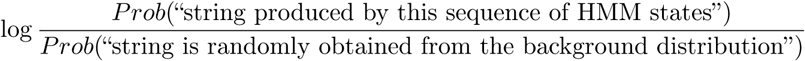

and is known as *log-odds score* [6].

For given HMM graph and background define the alignment score *Score*(𝓛) for the string 𝓛 = (*L*_1_*, …, L_n_*) as the best score along all its possible global alignments 𝓐 = (*E*_1_*, …, E_n_*).

We consider the following problem of graph alignment to profile HMM: for given sequence graph *G*, profile HMM *pHMM*, background distribution 𝓑, and integer *k >* 0, among all the paths in *G* obtain *k* different paths with the highest possible scores. We will call this problem *Top(k) path problem* throughout the article.

It is well known that for the one string the best score alignment could be found by Viterbi dynamic programming algorithm [18]. Here we seek for the generic solution of the alignment problem: (1) among all paths in the sequence graph, and (2) obtaining *k* best paths instead of just one.

### 2.2 Event Graph

We present the solution of *Top(k)* path problem via the special data structure called *event graph*.

First, define the expanded sequence graph 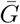 as graph *G* with added special Empty vertex with no letter on it. Connect Empty to all other vertices by bidirectional edges.

Each vertex (or *event*) of an event graph is a pair (*E, P*) of an emissive (I or M) HMM state *E* and a non-empty position *P* of 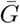 or one of two special vertices: Source = (Start, Empty) and Sink = (End, Empty).

A directed edge from event (*E*_1_*, P*_1_) to event (*E*_2_*, P*_2_) exists only if and only if: there is an edge from *P*_1_ to *P*_2_ in 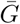; emitters are consequent, i.e., there is a path in HMM from *E*_1_ to *E*_2_, that does not contain other emitters. Due to the used pHMM structure there could be only one such path between the states.

The weight of vertex (*E, P*) equals to log(𝓔(*E, Letter*(*P*)))−log(𝓑(*Letter*(*P*))). Edge weight is a sum of logarithms of the probabilities of transitions on the edge. Since these probabilities are always less or equal to 1, edge weights are always non-positive, while vertices could have positive weights due to the presence of background distribution.

One can easily see that each path from Source to Sink corresponds to a global alignment of the pHMM against some path in *G* whereas path weight (sum of weights of edges and vertices along the path) is equal to the alignment score.

### 2.3 Top(1) Path Problem via Event Graph

Event graph allows one to construct the solution of Top(1) problem straight from the definition.

#### Algorithm 1: Top(1) path finding

**Input:** sequence graph *G*, profile HMM *pHMM*, background distribution *Background*

1 Construct event graph from *G*, *pHMM*, and *Background*

2 Find the best (highest scored) event path from Source to Sink

3 Extract sequence path (series of positions) and the correspondent string from the best event path

One could easily see that the problem of finding the best event path in general does not have a finite solution. Indeed, if the event graph is acyclic or at least has no cycles of positive weight, then the solution will certainly exist. In the general case event graph could have cycles due to the presence of I-loops in pHMM and depending to a particular background distribution some of these cycles may positive weight. If there is a path from Start to End that contains one of such cycles, then one can roll on it endlessly increasing the score and therefore the proper solution would not exist.

Fortunately, this problem is more theoretical than practical. In our implementation we use a simple heuristic to deal with it. We find all I-loops of positive weight in pHMM and unroll them into *C* (default: 10) sequential I states. This procedure is equivalent to restricting the total number of emissions. Typically there is only small fraction of such I-loops (not exceeding 2.5% for all considered HMMs and therefore the size of pHMM graph increases negligibly.

### 2.4 Dynamic Programming Algorithm for Best Path Extraction

For each vertex and each edge of the event graph find the best (having largest score) path from Source. Denote those best score values as *DIST* (vertex) and *DIST* (edge). Note that due to absence of positive-weighted cycles those *DIST* s always exist and are finite.

The values of *DIST* s could be calculated via the Bellman-Ford algorithm [4]. However, the computational complexity of this approach is 𝓞(|*V* (*EventGraph*)| × |*E*(*EventGraph*)|), that is completely impractical. Still, due to special layered event graph structure, *DIST* s could be computed significantly faster level-by-level. One possible approach to this is described in Algorithm 2.

The only non-straightforward step in Algorithm 2 is I-loop relaxation. Due to negativity of all I-loop edges we can sequentially update I-loop events *DIST* using a priority queue as in classical Dijkstra algorithm [4]. Overall algorithm complexity is (𝓞*|pHM M| × |E*(*G*)| log |*V* (*G*)|), where log comes from priority queue push/pop costs.

For sequence graphs |*E*(*G*)| ~ |*V* (*G*)| since vertex degrees are bounded by above (e.g. by 4 for de Bruijn graphs), therefore, we will count them together denoting *|G|* = *|V* (*G*)*|* + *|E*(*G*)*|*. Thus, the overall complexity is *𝓞*(*|pHM M* | × |*G* | log | *G*|)

Having *DIST* (Sink) calculated we can easily reconstruct the best path by backtracking: start from Sink vertex and go backwards into each vertex taking incoming edge with the highest *DIST* until we reach Source. *k* best paths could be found iteratively using the observation that the *i*-th shortest path in the sequence must branch from one of the *i* − 1 shortest paths already identified (so-called Eppstein algorithm, [7]).

### 2.5 From Top(1) to Top(*k*)

Though there is a simple solution for the extraction of the best *k* paths from the event graph, the problem of finding top sequence graph paths is still non-trivial and could not be easily solved by extracting *k* best paths from SOURCE to SINK.

The first problem here is that top *k* alignments are not equal to top *k* sequences. Indeed, full event graph represents all possible alignments of the pHMM against all sequence graph paths. Each sequence path has a combinatorial number of alignments against the pHMM. And each alignment corresponds to its own path in the event graph. The highest scored path in the full event graph indeed corresponds to the best alignment of the best sequence path. However, the second best alignment usually corresponds to the second best alignment of the same sequence path and not of some other sequence path.

#### Algorithm 2: Level-by-level *DIST* computation (optimized Bellman-Ford algorithm)

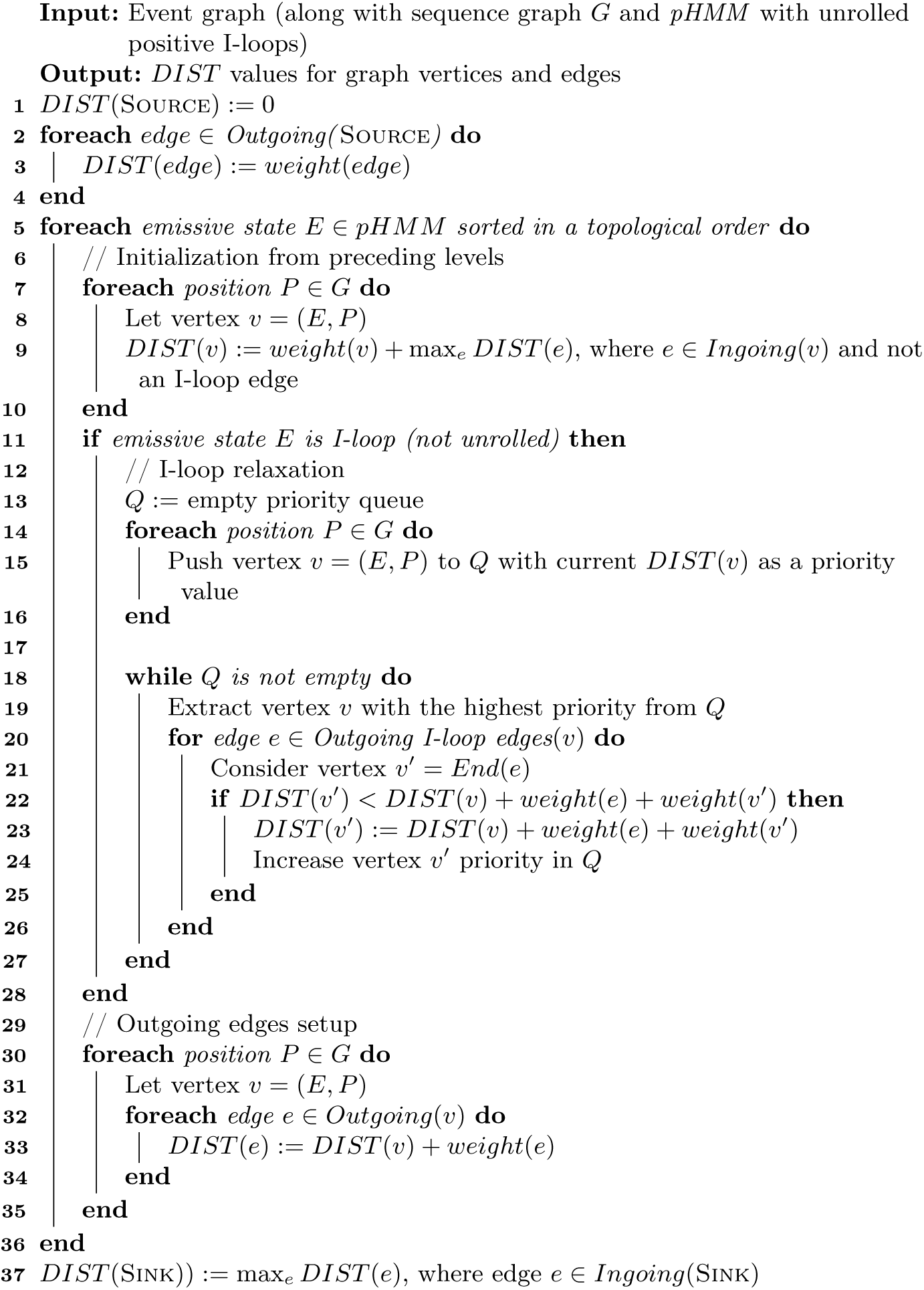

The second issue is driven by particular applications. Among top *k* paths we would like to see really different paths rather than paths that differ only by adding/trimming a small number of letters at the beginning/End. Typically extending/trimming the sequence in such way changes the final score only slightly and variations of the best path displace all other paths from the top list.

The third issue is that the number of different sequence paths with the proper score could also be combinatorial. For the string with d single-letter variations (corresponding to a path with *d* bubbles) the number of different paths would be 2^*d*^. All the paths would be slightly different and would probably have the same or almost the same score. Without additional information we could not prefer one path to another and report an answer of a reasonable (non-combinatorial) size.

This way the proper Top(*k*) problem statement might look like as follows: find *k* sufficiently (in the terms of the second issue) different paths in the initial sequence graph that maximize alignment score, but such problem seems to be very hard to solve. The only solution we know is increasing the number of extracted top alignments and further result filtering, but this approach is impractical for large *k* since the number of paths to extract is not known in advance. Also, the construction, storage and operations on full event graph (consisting of |*pHMM* | × |*G*| vertices) is time and memory consuming.

To deal with these issues we propose an heuristic solution that could significantly speed up Top(*k*) paths extraction making it possible even for very large *k* (say, *k >* 100000) that could be useful for complicated metagenome graphs where the number of candidate paths could be really large. This approach introduces a new structure representing alignment of the whole sequence graph against the pHMM that could be also interesting and useful by itself. Particularly, it helps to overcome the third issue, allowing one to consider relatively compact object rather than an endless number of slightly different paths.

### 2.6 Collapsing Event Graph

Let us consider a subgraph of the full event graph called collapsed and trimmed event graph.

*Collapsed event graph* is a subgraph of an event graph where for each vertex (*E, P*) (including Source and Sink) all its ingoing and outgoing neighbours (vertices linked by ingoing and outgoing edges respectfully) have different positions. This way different Start-to-End paths in collapsed event graph correspond to different paths in the initial sequence graph.

*Trimmed event graph* is a subgraph where each vertex does not have ingoing edge from SOURCE and ingoing edge(s) from non-SOURCE vertex at the same time. The same property is ensured for outgoing edges and Sink.

All the different Source-to-Sink paths in collapsed and trimmed event graph correspond to the paths in the initial sequence graph that are not substring or superstring of each other and do not have a perfect overlap, i.e. any prefix of one path cannot not be a suffix of another.

If we perform *k* best paths finding algorithm on collapsed and trimmed event graph, we extract *k* non-trivially different sequence paths. There are multiple ways how event graph could be trimmed and collapsed. Algorithms 3 and 4 make an arbitrary subgraph of an event graph collapsed and trimmed but still keeping the best scored paths intact.

#### Algorithm 3: Event graph collapsing

**Input:** Event graph or its arbitrary subgraph

1 For each vertex and each edge calculate the *DIST* (vertex) and *DIST* (edge) values using the same DP procedure as in best path finding algorithm (Section 2.4)

2 For each vertex, group all incoming edges by position of an incident vertex. In each group find the edge with the highest score and remove other edges from the graph

3 Do steps 1 and 2 on the graph backwards: with all the edges reversed and going from Sink to Source

#### Algorithm 4: Event graph trimming

**Input:** Event graph or its arbitrary subgraph

1 Compute *DIST* (edge) for each edge as in the previous algorithm

2 For each vertex incident to Source: remove all incoming edges worse than the incoming edge from Source. If there are still other incoming edges, remove the edge that goes from the Source vertex

3 Do steps 1 and 2 on the graph with all the edges reversed considering edges going to Sink

After collapsing and trimming we perform graph cleanup by removing all vertices unreachable from SOURCE or SINK.

#### Algorithm Discussion and Analysis

Both algorithms preserve the best path intact since for each vertex that belongs to the best path incoming and outgoing edges with maximum *DIST* value belong to the best path as well.

Collapsing procedure preserves existing sequence paths, but may reduce their scores. For each SOURCE-to-SINK path that got broken during the collapsing step there still exists another event path with the same corresponding sequence path. Note that collapsing removes edges, but it leaves one edge going to each position, in other words: for each sequence path *P*_1_*, …, P_n_* collapsing procedure will preserve at least one corresponding event path.

Collapsing procedure can reduce score of a path therefore yielding suboptimal alignment. If the sequence path shares prefix with another path with better score, the prefix alignment will be inherited from the other path. The same goes for the suffix. Our experiments show that this phenomenon is usually not a big problem for the real graphs. Even more, in order to address this problem one could re-align the extracted sequence paths against pHMM.

Trimming procedure could break existing sequence path but if it does so, then there does exist a path slightly different from this one with better score.

Full event graph represents all possible alignments of all graph paths against the pHMM. Collapsed and trimmed event graph represents an alignment (probably suboptimal, but still reasonable) of the whole graph against the pHMM. It could be viewed as an analogue of partial order graph alignment [10] but for graph paths rather than separate homologous sequences.

### 2.7 Alignment Algorithm: Collapsed Event Graph Construction

Collapsed trimmed event graph takes sufficiently less space and could be constructed on-fly, therefore we do not need to construct full graph and then collapse it. Algorithm 5 represents such an approach.

#### Algorithm 5: pHMM alignment against sequence graph

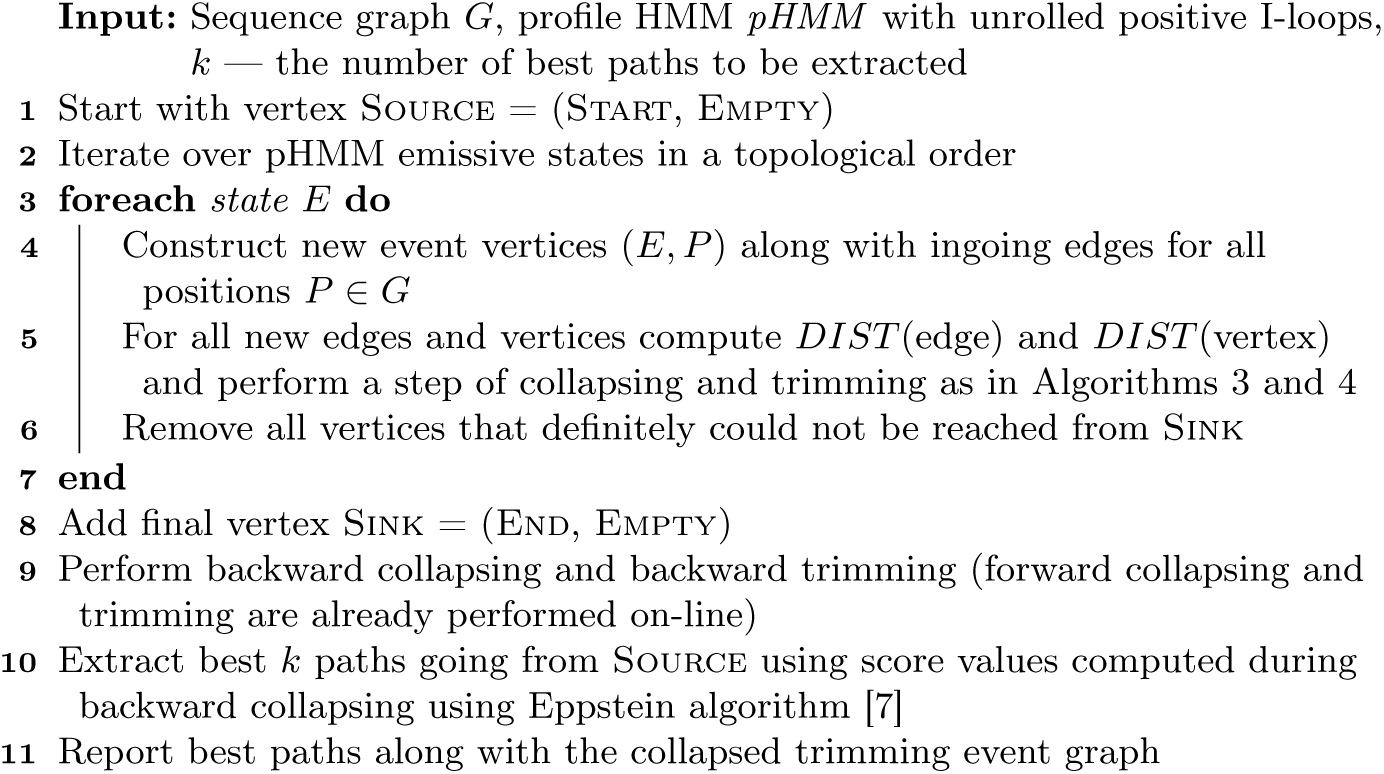

Two parts contribute to the algorithm complexity: collapsed event graph construction and extraction of best paths. The first part has the computational complexity of 𝓞(|*pHM M| × |G*|log|*G* |), the same as general Top(1) algorithm (see Section 2.4). The second part (Eppstein algorithm [7]) has complexity 𝓞(*kN* log(*kN*)), where *N* is average path length and log is from priority queue overhead. Note that in collapsed event graph incoming vertex degree is bounded like in sequence graph (usually by 4 for nucleotide graphs), therefore there is no additional term in the complexity equation.

### 2.8 Implementation and heuristics

We implemented the described approximate Top(*k*) algorithm solution in the PathRacer tool. PathRacer takes as input de Bruijn graph in GFA format (only SPAdes or SPAdes-compatible graphs are currently supported) and nucleotide pHMM. For amino acids HMMs we perform on-fly translation that allows us to consider the initial graph as a sequence graph over extended (20 amino acids and stop codon) amino acid alphabet.

The algorithm 5 has overall complexity (𝓞*|pHM M|×|G|*log|*G|*+*kN* log(*kN*)), however, the first component in the sum might be intractable for large graphs. In order reduce the computation complexity we implemented several heuristics:

1. We run hmmalign algorithm from HMMER 3 [6] on the edges of the input sequence graph and consider only the neighborhood of suitable size of matched edges: we walk forward and backward from each matched edge not far than the length of the corresponding pHMM overhang and then join all visited positions and extract sequence subgraph. HMMER options (e.g. domain E-value thresholds) are available to control the size of seed set.
2. During the event graph construction after each step new vertices are filtered out:
  – if the continuation of alignment through this vertex will always reach a dead-end before reaching Sink vertex;
  – if the continuation of alignment through this vertex will produce stop codon before reaching Sink vertex (for amino acid HMM);
  – all vertices with the score lower than a threshold parameter (default: −250);
  – all vertices except *T* top score vertices. The number *T* is reducing while moving along the pHMM.

This heuristic improves both the running time and memory consumption drastically (see Supplementary Table 1) with no significant changes in the results.

## 3 Results

In order to demonstrate the viability of our solution we conducted four experiments using the variety of datasets and pHMMs.

### 3.1 23S Search in *E. coli str. K12* Assembly

*E. coli* genome contains seven ribosomal RNA (rRNA) operons. Each operon contains a 16S rRNA gene, a 23S rRNA gene, and a 5S rRNA gene (except for one operon, which contains two 5S rRNA genes) interspersed with various tRNA genes.

We considered the *E. coli str. K12* dataset from [3]. The Illumina reads were of length 100 bp with mean insert length 270 bp. SPAdes 3.12 [13] assembly was performed in the normal multi-cell isolate mode with default settings and the resulting assembly graph in GFA format was obtained. Quick check of the results revealed that SPAdes assembled the 16S gene on the single contig, but was unable to derive the whole sequence of 23S gene. Probably the reason is that among seven copies of 23S gene, six were inexact, still 3 variants of 23S gene were scaterred over 3, 3 and 2 edges correspondingly. Other variants were absent in the assembly graph and probably were collapsed during the graph simplification procedures.

We aligned final assembly graph to prokaryotic rRNA pHMM models from [16] and all 3 variants of 23S gene were the top paths extracted by PathRacer (see Figure 2). No other (false) paths were produced.

**Fig. 2.**
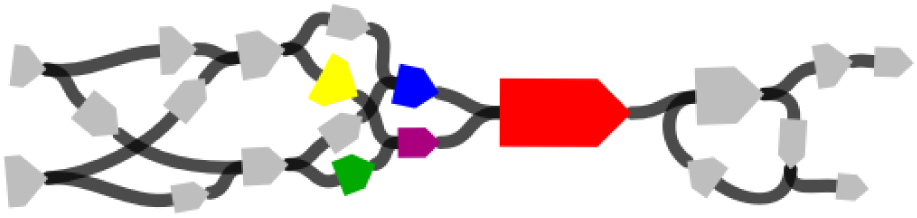
*E. coli str. K12*: extracted 23S paths (yellow-violet-red, green-violet-red and blue-red) and their neighborhood

### 3.2 16S Components of SYNTH Mock Metagenome Dataset

To showcase the abilitiy of PathRacer to deal with complex metagenomic data we considered 16S genes contained in SYNTH mock metagenome dataset from [17].

Synthetic community data set (SYNTH) is a set of reads from the genomic DNA mixture of 64 diverse bacterial and archaeal species (SRA acc. no. SRX200676) that was used for benchmarking the Omega assembler [9]. The dataset contains 109 million Illumina HiSeq 100-bp paired-end reads with mean insert size of 206 bp. The reference genomes for all 64 species forming the SYNTH dataset are known.

The assembly was performed by metaSPAdes 3.12 [13] with default parameters. To increase the sensitivity we considered *strain* (rather then the final *consensus*) assembly graph. We aligned the graph against 16S profile HMM from [16] and analyzed paths reported. PathRacer reported 1088577 paths. It is a reasonable number since the corresponding graph component is very complicated (see Figure 3) and therefore many chimeric alignments are expected. From SILVA [15] we obtained 179 16S sequences (most of the species have several variants of this gene), constructed BLAST database from all reported paths and performed a search in this database of all 16S gene sequences from SYNTH. For 55 out of 64 species all the 16S sequences were found with >95% BLAST identity. For 22 species at least one of 16S sequences was found with 100% BLAST identity (some variations were missing).

**Fig. 3.**
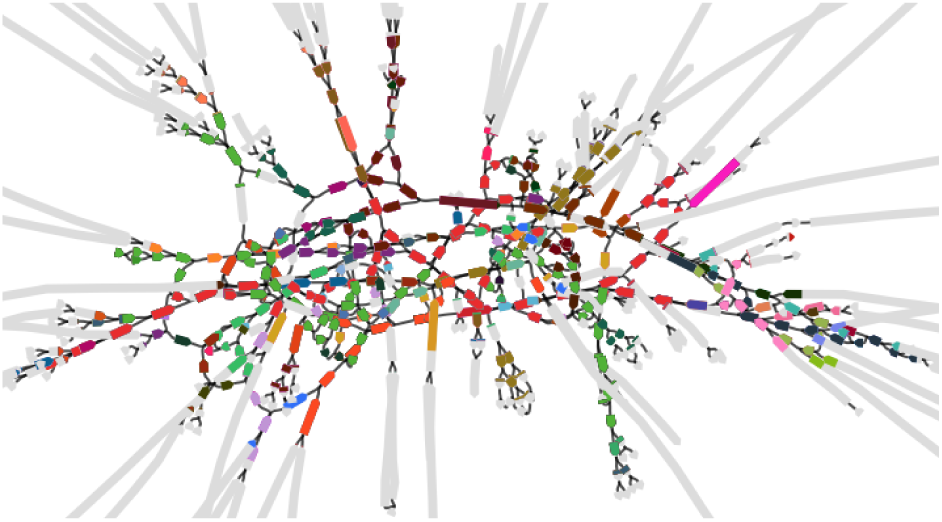
SYNTH: 16S matches of known sequences and their neighborhood. Different species are colored differently

### 3.3 Urban Wastewater Metagenome

In [12] comparative metagenomics was used to investigate the occurrence of antibiotic resistant genes in wastewater and urban surface water environments in Singapore. For this experiment we took H1 (clinical isolation ward, SRA acc. no SRR5997548) dataset and performed assembly by metaSPAdes 3.12 with default settings. As with SYNTH dataset we used *strain* assembly graph to increase the sensititivy of the search.

The assembly graph was aligned by PathRacer against selected betalactamase gene pHMMs obtained from NCBIfam-AMR database [1]: *bla*_CTX*−*M_, *bla*_IMP_ and *bla*_TEM_ among the others. The resulting paths consisting of several graph edges could be seen on Figures 4,5,6.

**Fig. 4.**
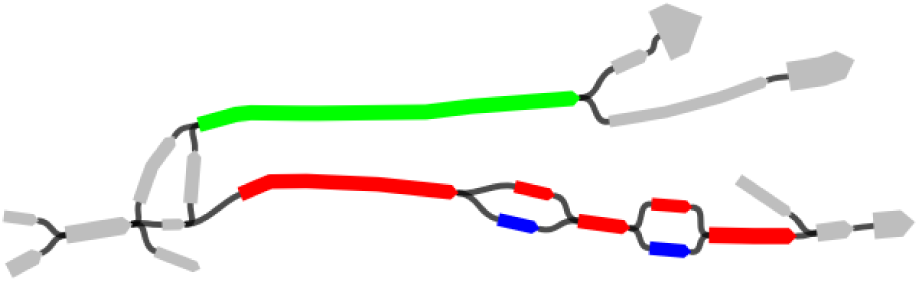
*bla*_CTX−M_ paths and their neighborhood. Green path corresponds to CTX-M-15 family; blue and red corresponds to CTX-M-9 and CTX-M-14 respectively

**Fig. 5.**
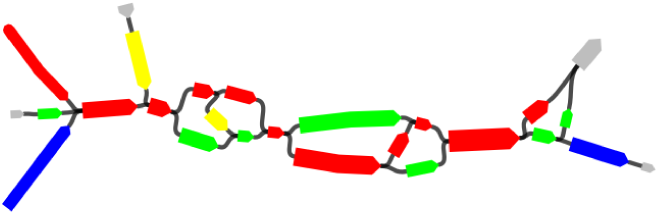
*bla*_IMP_ paths and their neighborhood. Red path corresponds to IMP-1 family

**Fig. 6.**
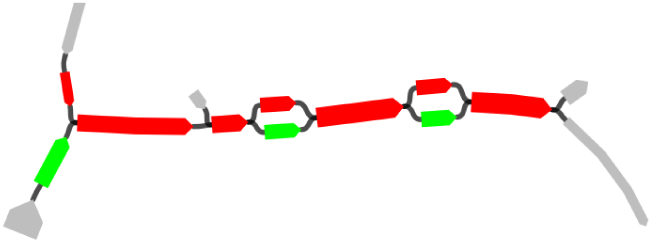
*bla*_TEM_ paths and their neighborhood. Red path corresponds to TEM-1 family

Different non-overlapping paths on the Figure 4 represents several different families of *bla*_CTX*−*M_ gene: green corresponds to CTX-M-15 family, while blue and red corresponds to CTX-M-9 and CTX-M-14 respectively (differs from each other only by 2 SNPs). Note that Table S5 in [12] mentiones the presence of CTX-M-15 and CTX-M-18 beta-lactamase families in this dataset, however we found that the extracted graph path that corresponds to CTX-M-9 aligned to the reference with 100% identity. Since the amino acid sequence of the CTX-M-18 beta-lactamase differs from that of the CTX-M-9 beta-lactamase by an Ala-to-Val change at position 231 [14], the additional checking of the results of [12] might be necessary (in [12] different gene sequences were searched in reads rather than assemblies and therefore might be prone to sequencing errors).

### 3.4 PathRacer and MegaGTA Running Time and Memory Consumption

We compared PathRacer performance with MegaGTA using 16 threads on urban dataset on all 159 beta-lactamase pHMMs obtained from NCBIfam-AMR database [1]. The memory consumption and running times are presented in Table 1.

**Table 1.** PathRacer and MegaGTA benchmark: running time and peak memory consumption. *urban* dataset, 159 beta-lactamase HMMs, 16 threads. PathRacer performance with disabled event graph online filtering is also shown

| Tool | Graph assembly | pHMM alignment |
| --- | --- | --- |
| MEGAGTA | 30m / 30GiB | 11h / 30GiB |
| METASPADES + PATHRACER | 1h50m / 20GiB | 50m / 7GiB |
| METASPADES + PATHRACER (no filtering) | 1h50m / 20GiB | 2h20m / 11GiB |

**Table 2.** PathRacer and MegaGTA results for *urban* data. blaCTX-M, blaIMP, blaTEM beta-lactamase gene families. Annotation performed by online NCBI BLAST [2], only results with 100% coverage and identity included

| Gene family | As shown in [12] | MEGAGTA | PATHRACER |
| --- | --- | --- | --- |
| *bla*<sub>CTX-M</sub> | blaCTX-M-15;<br>blaCTX-M-18 | OXY-1-1/2/3 (177aa);<br>CTX-M-15 (291aa);<br>CTX-M-3 (291aa);<br>CTX-M-9 (291aa);<br>SFO-1 (295aa);<br>CTX-M-14 (291aa);<br>+ 4 chimeric sequences<br>(247aa, 203aa, 295aa,<br>242aa) | CTX-M-15 (291aa);<br>CTX-M-14 (291aa);<br>CTX-M-9 (291aa);<br>SFO-1 (295aa) |
| *bla*<sub>IMP</sub> | blaIMP-1 | IMP-1 precursor<br>+ 8 chimeric sequences<br>(245aa × 8) | IMP-1 precursor |
| *bla*<sub>TEM</sub> | blaTEM;<br>blaTEM-157;<br>blaTEM-163;<br>blaTEM-211 | TEM-1 (286aa);<br>+ 43 chimeric and frag-<br>mented sequences | TEM-1 (286aa) |
| *bla*<sub>KPC</sub> | blaKPC2 | KPC-2 (293aa);<br>+ 113 chimeric and frag-<br>mented sequences | KPC-2 (293aa) |

We need to note that along with the input profile HMM files MegaGTA also requires original gene sequences and a set of gene sequences for FrameBot tool [20] that is used inside Xander/MegaGTA pipeline. Contrary to this, PathRacer needs input graph and pHMM file only. We have not included Xander into the benchmarking since MegaGTA is its improved version and it outperforms Xander as it shown in [11].

## 4 Conclusion

PathRacer utilizes both the assembly graph topology and the information about the known genes encoded in profile HMM during its operation. This way the putative gene sequences could be reconstructed even from the fragmented metagenomic assemblies; it does not matter whether the gene sequence is located within the single contig or scattered across several edges of an assembly graph.

Currently PathRacer could be viewed as a standalone tool that might be integrated into analysis pipeline. Though we anticipate that deeper integration into assembler pipeline might be possible, e.g. one could utilize the information from paired-end reads to filter out more paths in the even graph, or, use pHMM alignment to supplement repeat resolution and scaffolding.

## Acknowledgments

This work was supported by the Saint Petersburg State University (grant number 15.61.951.2015). The authors would like to extend a special thanks to Sergey Nurk and Tatiana Dvorkina for all the fruitful discussions that were of great help in improving the algorithms.

1 So far only only GFA from de Bruijn graph assemblers like SPAdes and MegaHit is supported, but we will address this restriction in the next PathRacer versions

